# Transdiagnostic Phenotyping Reveals a Host of Metacognitive Deficits Implicated in Compulsivity

**DOI:** 10.1101/664003

**Authors:** Tricia X.F. Seow, Claire M. Gillan

## Abstract

Recent work suggests that obsessive-compulsive disorder (OCD) patients have a breakdown in the relationship between explicit beliefs (i.e. confidence about states) and updates to behaviour. The precise computations underlying this disconnection are unclear because case-control and transdiagnostic studies yield conflicting results. Here, a large general population sample (N = 437) completed a predictive inference task previously studied in the context of OCD. We tested if confidence, and its relationship to action and environmental evidence, were specifically associated with self-reported OCD symptoms or common to an array of psychiatric symptoms. We then investigated if a transdiagnostic approach would reveal a stronger and more specific match between metacognitive deficits and clinical phenotypes. Consistent with prior case-control work, we found that decreases in action-confidence coupling were associated with OCD symptoms, but also 5/8 of the other clinical phenotypes tested (8/8 with no correction applied). This non-specific pattern was explained by a single transdiagnostic symptom dimension characterized by compulsivity that was linked to inflated confidence and several deficits in utilizing evidence to update confidence. These data highlight the importance of metacognitive deficits for our understanding of compulsivity and underscore how transdiagnostic methods may prove a more powerful alternative over studies examining single disorders.

## Introduction

Intentional decisions are dependent on the interplay between behaviour and beliefs. Beliefs guide behaviour, and the consequences of our behaviour in turn update beliefs. Computational models of learning suggest that the strength of belief (i.e. “confidence”) governs the extent of its influence on action; the less confident we are, the less our behaviour is influenced by pre-existing beliefs, compared to new information^1,2^. A breakdown in the relationship between action and belief is suggested to be characteristic of compulsive behaviours, e.g. in obsessive-compulsive disorder (OCD) or addiction. In these disorders, behaviour often appear autonomous, unguided by conscious control or ‘ego-dystonic’, such as persistent drug use despite negative consequences^3^ or out-of-control repetitive checking despite knowing the door is locked^4^. One potential cause of the divergence between intention and action in compulsive individuals is an impairment in the brain’s goal-directed system, which links actions to consequences and protects against overreliance on rigid habits^5^. Goal-directed planning deficits have been consistently observed in OCD^6–9^ and related disorders^10^ - there is evidence to suggest this constitutes a transdiagnostic psychiatric trait linked to several aspects of clinically-relevant compulsive behaviour.

Despite this, the precise mechanism supporting this dysfunction is only partially understood as most employed tasks struggle to separate the construction of an internal model (e.g. action-outcome knowledge) from its implementation in behaviour. Those that have attempted this have yielded interesting, if equivocal, results. One study showed that OCD patients get stuck in habits even when they possess the requisite action-outcome knowledge to theoretically perform in a goal-directed fashion^7^. This suggests that the implementation of goal-directed behaviour is deficient in OCD, independent of their ability to construct the model. However, this does not mean the internal model is intact; studies using more challenging tasks have found deficits in the acquisition of explicit action-outcome contingency knowledge itself in OCD patients^9^, suggesting that patients may have problems with both. These findings come from paradigms where instrumental action typically affects the kind of information that is gathered and thus are somewhat confounded and difficult to interpret. Recently Vaghi and colleagues addressed this metacognitive question in OCD patients with more precision using a paradigm that examined how patients make trial-wise adjustments to behaviour (i.e. implicit model) and confidence (i.e. explicit model) in response to feedback^11,12^. They found that in OCD, the association between confidence and behavioural updating (‘action-confidence coupling’) was diminished - patients’ behaviour did not align with their internal model. Further, while confidence estimates did not differ from healthy controls, OCD patients showed abnormalities in their learning rate, making more trial-wise adjustments in response to feedback than controls^12^.

The finding of intact confidence in OCD is consistent with prior work in perceptual decision-making where individuals high versus low in OCD symptoms had no differences in their confidence^13^ — results echoed by two large internet-based samples (N > 490) with the same task that also found no relationship to OCD symptoms^14^. A problem with this type of study design, however, is that it fails to capture the potentially competing influence of co-occurring disorders/symptoms in psychiatric populations. Even in studies where certain co-morbid diagnoses are explicitly excluded for, as in Vaghi *et al.*, rates of depression and anxiety are greater than controls^12^. Similarly, when self-report anxiety and depression severity are matched across groups by design, as in Hauser *et al.*^13^, this does not accurately reflect the average OCD patient where co-morbidity is the rule, not the exception (e.g. >25% of OCD patients are co-morbid for ≥4 additional diagnoses^15^). Indeed, selecting for individuals with high OCD scores but low depression scores might reflect statistical anomalies rather than a true and lasting absence of depression in those individuals. An alternative approach measures these relevant co-occurring symptoms in the same individuals and seeks to account for their (competing or inflating) influence on the cognitive measure of interest. We took this approach in a prior study and found that confidence abnormalities (which were not reliably linked to OCD or depressive symptoms in the same sample) were robustly associated with two transdiagnostic psychiatric dimensions in *opposing* directions: ‘anxious-depression’ was associated with reduced confidence, while ‘compulsive behaviour and intrusive thought’ was linked to inflated confidence. Given this, it is possible that in prior case-control studies in OCD, metacognitive abnormalities are obscured by the competing influence of co-occurring depression.

To test this, here we used the same transdiagnostic methodology on an online sample of 437 participants who completed the same task from Vaghi and colleagues^12^. We investigated the extent to which trial-wise action adjustments were disconnected from confidence reports with self-reported OCD symptoms, and whether this action-confidence decoupling is specific to OCD or also manifested in other psychiatric symptoms. We then tested if transdiagnostic phenotyping would reveal a more specific result – that only the compulsive dimension (as opposed to anxious-depression and social withdrawal) would be related to the decoupling of confidence and behaviour. Lastly, we investigated if the decoupling arose from failures in action-updating or confidence, and, with a reduced Bayesian model^2,11,12,16^, explored if there were abnormalities in the way compulsive individuals used information (e.g. recent outcomes, unexpected outcomes, environmental uncertainty and positive feedback) to update these behavioural measures.

## Results

Participants (N = 437) performed a predictive-inference task in which their goal was to catch a flying particle by placing a bucket in the correct location (Figure S1). On each trial, they repositioned the bucket and rated how confident they were in catching the particle. After the behavioural task, participants completed an IQ test and a battery of self-report questionnaires assessing a range of psychiatric symptoms. Individual item-level responses on these questionnaires were transformed into three transdiagnostic dimensions using weights defined in prior study^17^: ‘anxious-depression’ (AD), ‘compulsive behaviours and intrusive thought’ (CIT) and ‘social withdrawal’ (SW).

### Action-confidence decoupling is linked to various psychiatric symptoms

In line with prior research, size of action updates (bucket position difference from trial *t* and *t*+1) were strongly related to confidence within-subjects (*β* = −8.85, *Standard Error (SE)* = 0.11, 95% Confidence Interval (CI) [−9.45, −8.25], *p* < 0.001) (Figure 1a), such that lower confidence was linked to larger updates, in the sample as a whole. Previous work by Vaghi *et al.* found that OCD patients exhibited reduced coupling between action and confidence compared to controls, which was correlated to the severity of self-reported OCD symptomology within the patient sample^12^. We tested the latter in a general population sample and replicated this result; OCD symptom severity was associated with significantly lower action-confidence coupling (*β* = 1.30, *SE* = 0.21, 95% CI [0.89, 1.71], *p* < 0.001, corrected) (Figure 1b). However, we found that this relationship was profoundly non-specific – all nine psychiatric symptoms tested showed a similar pattern of reduced coupling. 6/9 questionnaires (alcohol addiction, depression, eating disorders, impulsivity, OCD and schizotypy) had significant decoupling at *p* < 0.001 corrected; the remaining three (apathy, social anxiety, trait anxiety) trended in the same direction, but did not survive Bonferroni correction for multiple comparisons.

**Figure 1.**
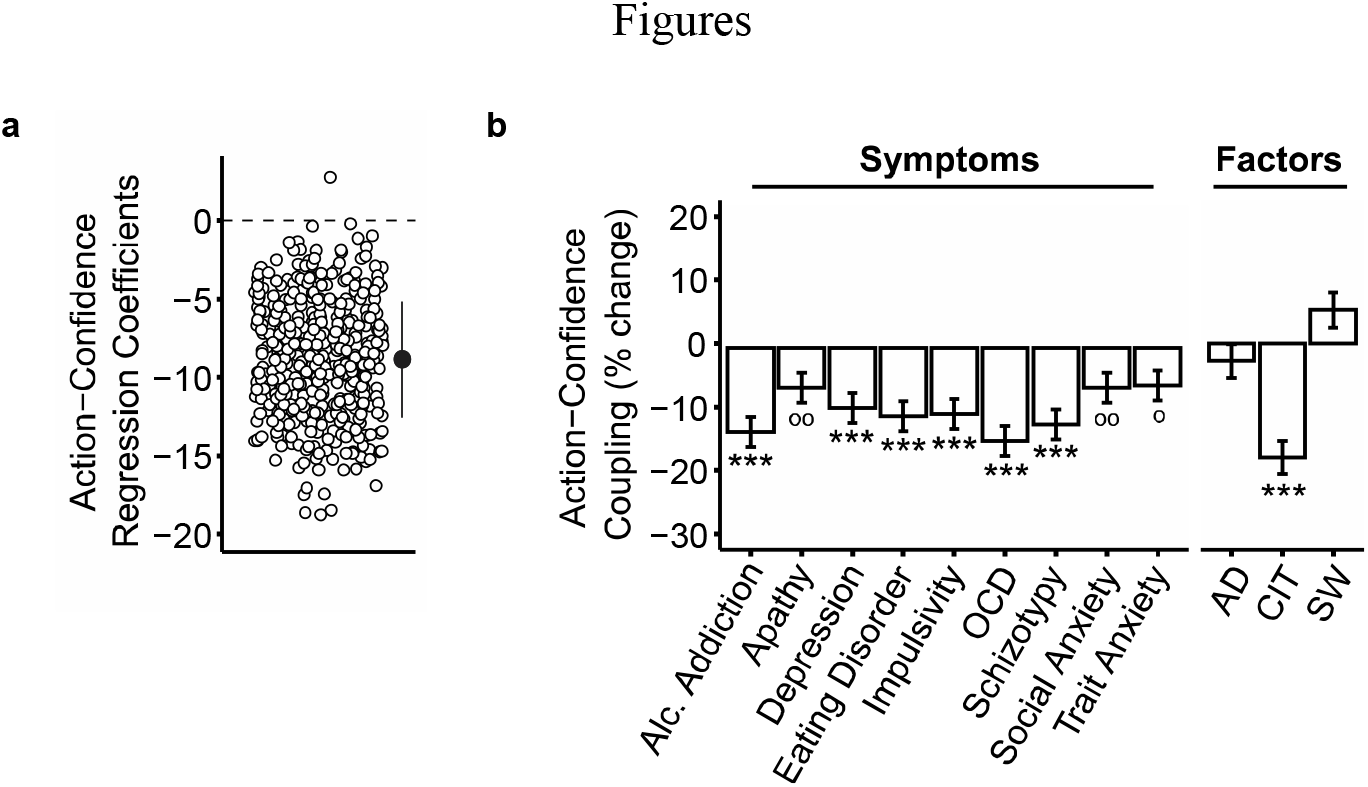
Action-confidence coupling and its relationship with psychiatric symptoms and factors (controlled for IQ, age and gender). AD: anxious-depression, CIT: compulsive behaviour and intrusive thought, SW: social withdrawal. (a) Regression model where action update is predicted by confidence. Individual coefficients are represented by circles. Marker indicates the mean and standard deviation. As expected, regression coefficients were negative, such that higher confidence was associated with smaller updates to the bucket position (‘action’). (b) Associations between action-confidence coupling and self-reported psychopathology or psychiatric factors. All symptoms predicted a decrease in action-confidence coupling. This decoupling relationship was specifically captured by the CIT dimension with its effect size larger than for any individual questionnaire. Each psychiatric symptom was examined in a separate regression, while factors were included in the same regression model. The Y-axes shows the percentage decrease in the size of the action-confidence coupling effect as a function of 1 standard deviation increase of symptom/factor scores. Error bars denote standard errors. °p < 0.05, °°p < 0.01 uncorrected, *p < 0.05, **p < 0.01, ***p < 0.001. Results are Bonferroni corrected for multiple comparisons over number of symptoms/factors. See also Figure S9.

### Transdiagnostic analysis shows a more specific pattern

When we refactored the data into three transdiagnostic dimensions defined previously in the literature, a profoundly different picture emerged. CIT was the only dimension to show decreased action-confidence coupling (*β* = 1.57, *SE* = 0.23, 95% CI [1.13, 2.01], *p* < 0.001, corrected) (Figure 1b). Thus, while reductions in action-confidence coupling show broad and non-specific relationships to all psychiatric symptoms studied here, this pattern is explained by a transdiagnostic dimension.

### Compulsivity is linked to inflated confidence, not aberrant action-updating

Prior work in diagnosed patients found no confidence biases in OCD, but abnormalities in action-updating. Using our transdiagnostic method, we found a strikingly different pattern of results. CIT was associated with higher overall confidence levels (*β* = 6.74, *SE* = 1.02, 95% CI [4.75, 8.73], *p* < 0.001, corrected), and not changes in action-updating. In line with prior work, we found that AD was associated with lower confidence (*β* = −3.42, *SE* = 1.04, 95% CI [−5.45, −1.39], *p* = 0.003, corrected) (Figure 2a). Because OCD patients tend to have high levels of AD, this finding suggests that a transdiagnostic method is necessary to reveal the role confidence plays in clinical phenotypes, which is otherwise obscured within the heterogeneous diagnostic category. In terms of action-updating, only SW showed an association, such that participants scoring high in this dimension tended to move the bucket more (*β* = 0.89, *SE* = 0.28, 95% CI [0.34, 1.45], *p* = 0.005, corrected) (Figure 2b).

**Figure 2.**
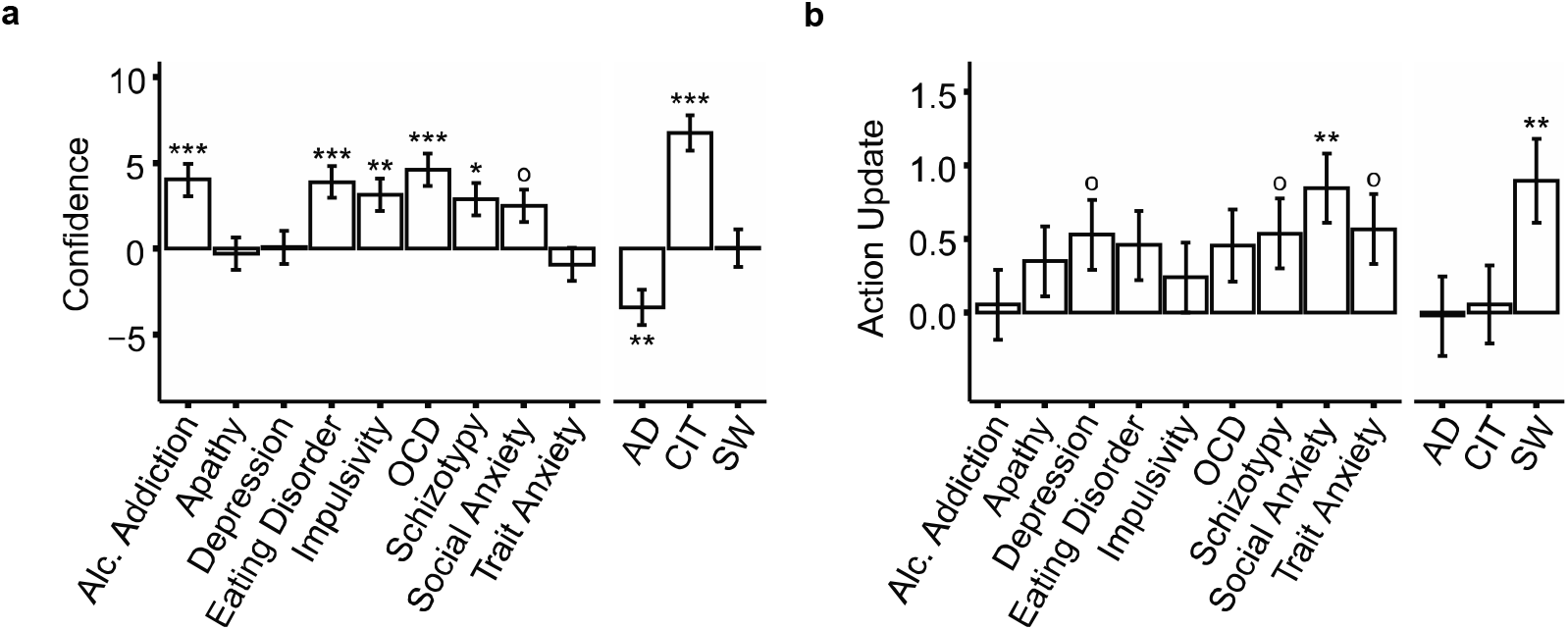
Associations between psychiatric symptoms, or transdiagnostic factors (controlled for IQ, age and gender) with confidence or action. AD: anxious-depression, CIT: compulsive behavior and intrusive thought, SW: social withdrawal. *(a)* Associations with confidence rating on each trial. Most of the questionnaires scores were positively associated with confidence. However, refactoring into transdiagnostic traits revealed previously obscured bidirectional associations. AD was linked to decreased confidence, while CIT was linked to increased confidence. *(b)* Associations with action updates (i.e. bucket movement from one trial to the next). Only social anxiety was associated with an increased tendency to move the bucket, and this was similarly captured by, and specific to, the SW factor. The Y-axes shows the percentage decrease in the size of the action-confidence coupling effect as a function of 1 standard deviation increase of symptom/factor scores. Error bars denote standard errors. °*p* < 0.05, °°*p* < 0.01 uncorrected, \**p* < 0.05, \*\**p* < 0.01, \*\*\**p* < 0.001. Results are Bonferroni corrected for multiple comparisons over number of symptoms/factors.

### Confidence in compulsivity is less sensitive to unexpected outcomes, environment uncertainty and positive feedback

The previous analyses suggest that confidence in compulsive individuals is both inflated and decoupled to behaviour. To understand the mechanism behind this, we tested the extent to which confidence estimates were sensitive to multiple factors that should drive belief-updating. Specifically, prior work has shown that trial-wise adjustments in behaviour are influenced by 1) information gained from the most recent change in particle location, 2) surprising large particle location changes owing to change-points and 3) uncertainty of one’s belief about the particle landing location distribution mean^16^. To separate the contributions of these factors, we computed three normative parameters with a quasi-optimal Bayesian model^2,11,12,16^ (see Supplemental Methods) to the sequence of particle locations experienced by each participant. The parameters of the model included PE^b^ (model prediction error, the tendency to update towards the most recent particle landing location), CPP (change-point probability, the likelihood that a surprising outcome had occurred) and RU (relative uncertainty owing to the imprecise estimation of the distribution mean based on previous outcomes).

We analysed trial-wise confidence using regression models with these parameters including a categorial Hit regressor (previous trial was a hit or miss), and controlled for age, gender and IQ. Overall, confidence was influenced by PE^b^, CPP, RU and Hit (Table S1). The CIT symptom dimension was associated with a significantly diminished influence of CPP (*β* = 0.05, *SE* = 0.01, 95% CI [0.03, 0.08], *p* < 0.001, corrected), RU (*β* = 0.05, *SE* = 0.01, 95% CI [0.03, 0.07], *p* < 0.001, corrected) and Hit (*β* = −0.03, *SE* = 0.01, 95% CI [−0.05, −0.01], *p* = 0.003, corrected) on confidence (Figure 3a and Figure S6). In other words, confidence estimates in CIT were less sensitive to unexpected outcomes, the uncertainty of the true distribution mean and whether the previous particle was caught (i.e. correct trial). These results suggest that confidence in highly compulsive individuals is not only inflated, it is also disconnected to several sources of environmental evidence. Interestingly, the failures in utilizing evidence do not explain away overall inflated confidence observed in CIT (*β* = 6.78, *SE* = 1.02, 95% CI [4.79, 8.76], p < 0.001, corrected), suggesting these might be distinct phenomena. There were no associations between AD or SW and the extent to which evidence influenced confidence (Figure S6).

**Figure 3.**
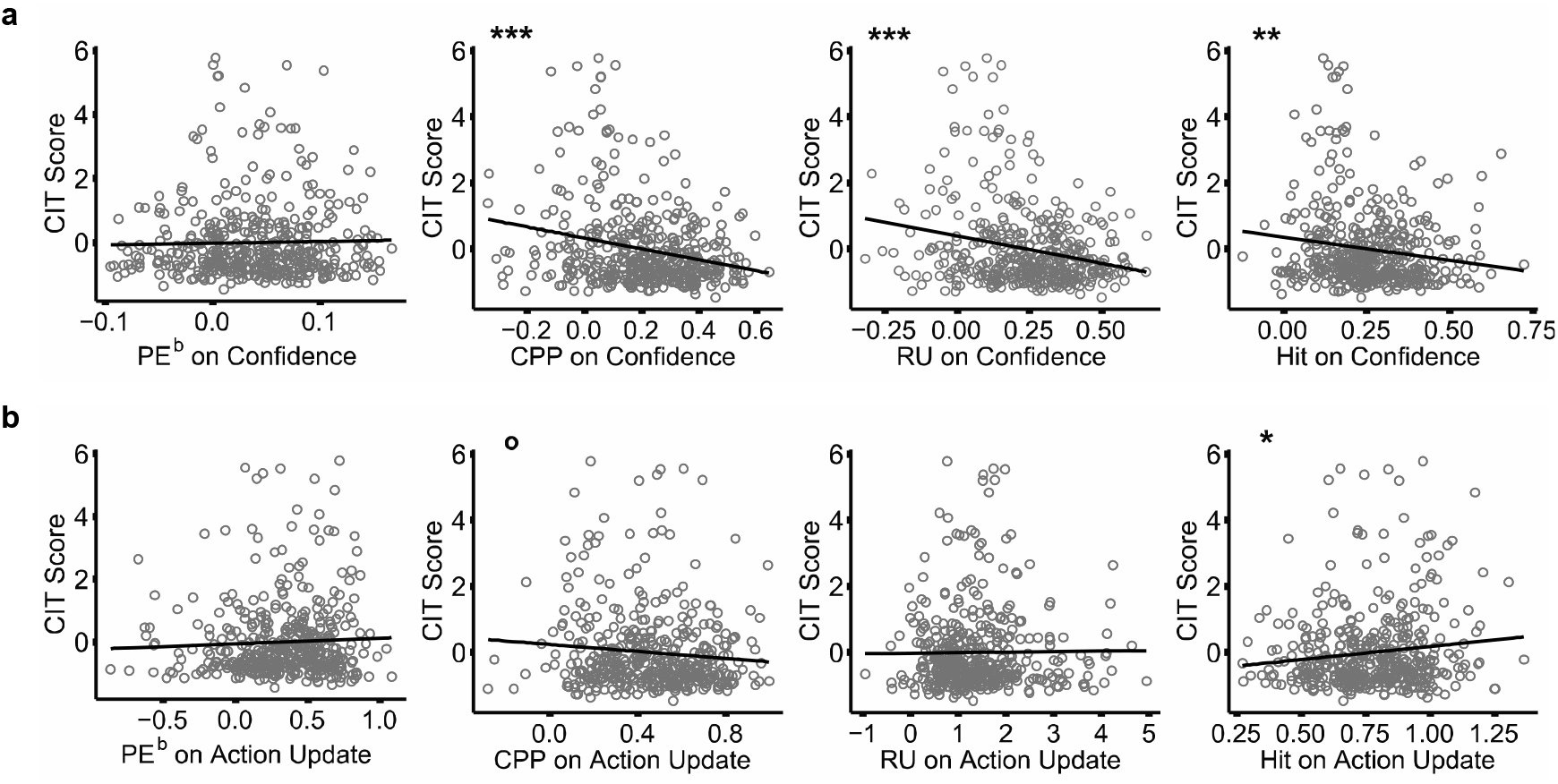
Confidence level/action update was predicted by absolute model prediction error (PE^b^), change-point probability (CPP), relative uncertainty (RU) and hit/miss categorial regressor (Hit), controlled for IQ, age and gender. Coefficient estimates from the model were correlated with ‘compulsive behavior and intrusive thought’ (CIT) severity. *(a)* CIT was found to be associated with significantly diminished influence of CPP, RU and Hit on z-scored confidence. PE^b^, CPP and RU on confidence coefficients are inverted to illustrate direction of effects. PE^b^: *r* = 0.02, 95% CI [−0.07, 0.12], *p* = 1.00; CPP: *r* = −0.24, 95% CI [−0.33, −0.15], *p* < 0.001; RU: *r* = −0.24, 95% CI [−0.32, −0.15], *p* < 0.001; Hit: *r* = −0.15, 95% CI [−0.24, −0.06], *p* = 0.005. *(b)* In contrast, CIT was found not linked to changes in the influence of any of model parameters on action update. For plotting purposes, we show the association of parameter and compulsivity without controlling for AD and SW. PE^b^: *r* = 0.05, 95% CI [−0.05, 0.14], *p* = 0.98; CPP: *r* = −0.10, 95% CI [−0.19, −0.003], *p* = 0.13; RU: *r* = 0.01, 95% CI [−0.08, 0.10], *p* = 1.00; Hit: *r* = 0.12, 95% CI [0.03, 0.21], *p* = 0.03. Note that a modest association with Hit on action update is observed here and illustrated in the associated plot but does not survive inclusion of all three factors in the same model. Circles represent coefficients of individual participants for model parameters from a basic model of confidence/action update ~ regressors*demographics (x-axis), against their CIT score (y-axis) (see Methods). Hit on action update coefficients are inverted to illustrate direction of effects, such that CIT is linked to an increase influence of hits on action-updating (which is negative in direction). CI = Confidence interval. °p < 0.05, uncorrected, *p < 0.05, **p < 0.01, ***p < 0.001. Results are Bonferroni corrected for multiple comparisons over the three factors. See also Figure S6.

### Action updates in compulsivity respond appropriately to evidence

Using the same approach described for confidence, trial-wise action adjustments were influenced by all model parameters (Table S1). In contrast to confidence, CIT was not linked to changes in the influence of any of the parameters on action (Figure 3b and Figure S6). SW was related to a significant increased influence of PE^b^, suggesting that individuals high in this trait had an increased tendency to update their action with every new outcome (*β* = 0.06, *SE* = 0.02, 95% CI [0.02, 0.09], *p* < 0.05, corrected) (Figure S6). There were no associations with AD. Additional analyses in the Supplement show that when demographics are not controlled for, some apparent associations between action-updating and compulsivity emerge that correspond to those reported previously in OCD^12^ (Figure S8).

## Discussion

In this study, we demonstrated that a breakdown in the relationship between explicit belief (confidence) and behaviour is associated with a transdiagnostic psychiatric dimension— compulsive behaviour and intrusive thought (CIT). This decoupling arises from abnormalities in belief, rather than behaviour. Individuals high in CIT exhibited overall inflated confidence estimates and failures in utilizing unexpected outcomes, belief uncertainty and positive feedback to inform their confidence levels appropriately. In contrast, action-updating in response to these factors did not differ as a function of severity of CIT. Our findings implicate a dysfunctional metacognitive mechanism in compulsivity that implicates difficulty in updating the explicit model of the world in response to various sources of evidence.

Existing models of compulsivity propose that deficits in goal-directed control leave individuals vulnerable to establishing compulsive habits^18^. Supporting evidence primarily come from behavioural tests, where OCD patients (and other compulsive disorders) have difficulty exerting control over well-trained habits when motivations change (i.e. a devaluation test)^6,7,9,19^. Other tasks have shown that compulsive patients have deficits utilizing a world model to make choices prospectively (even when habits are not present), relying instead solely on reinforcement (i.e. feedback) to direct choice^8,10^. Our current finding, that high compulsive individuals fail to update their world-model in response to several types of evidence, is an important extension of this literature. The challenge facing compulsive individuals has until now been presumed to be the implementation of the model rather than its generation and/or maintenance^18^. This implicates not just our understanding of compulsive disorders, but also their treatment. Recent work has shown metacognitive skills can be improved though adaptive training^20^; there is potential in these treatments for psychiatric populations where compulsivity is an issue.

Confidence was not just unresponsive to various factors underlying learning, it was also inflated in compulsive individuals. This finding replicates prior work that examined confidence in the context of perceptual decision making^14^, showing it extends to reinforcement learning – which is of course highly relevant for the behavioural aspects of compulsivity. Future work will be needed to dissect the specific mechanism through which confidence becomes inflated in compulsivity. In instances when confidence diverges from action, prior work has suggested confidence estimates may be corrupted by noise, internal states or a continued/lack of evidence processing^21,22^. Coupled with the finding that confidence is less informed by several sources of evidence in high compulsive individuals, it is possible that inflated confidence in compulsivity arises through some unmodeled source of information or noise.

This posit is supported by the finding that actions were updated normally in response to feedback in high compulsives, which accords with prior work showing that basic reinforcement learning in compulsive patient groups (i.e. ‘model-free’ learning) is intact^10,17^. That said, a previous study using this task found increased action-updating tendencies in OCD^12^. Here, the discrepancy does not appear to be explained by the superiority of a transdiagnostic approach per se, but our ability to control for some demographic confounds (see Supplemental Information, Figure S8). Instead, we found that social withdrawal (SW) was associated with a higher sensitivity to new information affecting action. Though this result was not hypothesized, we suggest that this might echo recent work where SW was found to be related to excessive deliberative processing^23^, a behaviour characteristic of socially anxious individuals where they excessively ruminate about previous social interactions, especially “if only” thoughts, for future social interactions^24^.

Beyond the specific results of this study with respect to confidence and compulsivity, our data highlight the benefit of transdiagnostic dimensions over traditional modes of phenotyping. When we examined questionnaires that are ubiquitous but rarely compared to one another in clinical research, we found strikingly non-specific patterns of association with task variables. For example, all nine questionnaires showed an association with action-confidence coupling in the same direction (6/9 surviving strict correction). In contrast, the compulsive factor was the only transdiagnostic dimension to show an association. In addition to resolving issues with collinearity across questionnaires, this approach also resolves issues associated with the heterogeneity within them. For example, severity of neither depression nor anxiety was associated with decreases in confidence using a standard clinical questionnaire, but the anxious-depression (AD) dimension was. In comparison to work with diagnosed patients, the benefits of the transdiagnostic approach are the same. Prior work using this task found no difference in OCD patients’ mean confidence ratings compared to healthy controls^12^, while we found a strong a reproducible association between CIT and inflated confidence and AD and diminished confidence^14^. Given that OCD is frequently co-morbid with anxiety disorders (over 75%^25^), which has an opposing relationship to confidence, it is no surprise that differences between OCD patients and controls are obscured when transdiagnostic dimensions are not considered. Together, these data suggest that transdiagnostic phenotyping may, at least in some domains, provide a closer fit to underlying brain processes than DSM distinctions.

This study was not without limitation. Our study was conducted online, thus experimenter control of the testing environment was virtually non-existent. Additionally, as the task was adapted for web-based testing, bucket movements were controlled by keyboard presses and not a rotor controller as in Vaghi *et al.*^12^, which may contribute to increased noise in spatial update measure. However, our ability to collect a large sample was evidently sufficient to mitigate the issue of increased noise in data collected online. We were able to reproduce basic main effects on model parameters similar to Vaghi *et al.*^12^ and also replicated previously observed associations with confidence, CIT and AD. With respect to the psychiatric dimensions, we not only reproduced the factor structure from a prior paper with our current data (Figure S7), we used the factor weights from this prior publication^17^ to transform raw questionnaire scores into transdiagnostic factors for analysis. This ensured independence and underscores the robustness and reproducibility of these factors and their association to cognition. The extent to which these results are applicable to diagnosed patients is not something we can directly address here. However, it is notable that we replicated the association between OCD symptoms and action-confidence decoupling observed in a clinical sample that were tested in-person^12^. The same applies to goal-directed planning, which is both deficient in patients tested in-person^9^ and correlated with OCD symptoms in the general population tested online^17,26^. Notably, recent work in a patient sample even found that goal-directed deficits were more strongly associated with the compulsivity dimension than OCD diagnosis status^27^. As such, there is no reason to suspect these findings are not applicable to patients. This method has several strengths over the case-control approach: it directly addresses the issue of psychiatric co-morbidity, helps us to achieve higher statistical power and thus promotes reproducibility, and makes research faster, more efficient and more representative^28^.

To conclude, we highlighted how a transdiagnostic methodology can be crucial for uncovering specific associations between pathophysiology and clinical symptoms. We used this method to show that compulsive behaviour and intrusive thought is associated with reduced action-confidence coupling, inflated confidence and diminished influence of evidence on confidence estimates. Our findings suggest that compulsivity is linked to problems in developing an explicit and accurate model of the decision space, and this might contribute to broader class of problems these individuals face with goal-directed planning and execution.

## Methods

### Power Estimation

Previous research utilizing the predictive-inference task were constrained to small sample psychiatric populations^12^. As such, we referred to earlier work that investigated confidence abnormalities in large general population cohorts with transdiagnostic symptom dimensions to determine an appropriate sample size^14^. The prior study reported an association of the ‘anxious-depression’ factor with lowered confidence level (*β* = −0.20, *p* < 0.001), an effect size suggesting that N = 295 participants were required to achieve 90% power at 0.001 significance level.

### Participants

Data were collected online using Amazon’s Mechanical Turk (N = 589). Participants were ≥18 years, based in USA and had >95% of their previous tasks on the platform approved. After reading the study information and consent pages, they provided informed consent by clicking the ‘I give my consent’ button. Participants were paid a base sum of 7 USD plus up to 1 USD bonus. Of the sample, 249 were female (42.3%) with ages ranging from 20-65 (mean = 36.3. SD = 10.2) years. All study procedures were approved by and carried out in accordance with regulations and guidelines of Trinity College Dublin School of Psychology Research Ethics Committee.

### Exclusion criteria

Several pre-defined exclusion criteria were applied to ensure data quality (see Supplemental Methods for details). In total, 153 participants (25.9%) were excluded, a rate typical for web-based experiments leaving 437 participants for analysis.

### Predictive inference task

We adapted the predictive-inference task from Vaghi *et al.* for web-based testing (Figure S1). Left and right arrow keys enabled response navigation while a spacebar press was used for decision confirmation. The task consisted of a particle flying from the centre of a large circle to its edge. First, participants positioned a ‘bucket’ (a free-moving arc on the circle edge) in order to catch the particle. Once bucket location was chosen, a confidence bar scaling 1 to 100 would appear below the circle after 500 ms. The confidence indicator would begin randomly at either 25 or 75. Participants then indicated how confident they were the particle would land in the bucket. After confirmation of the confidence report, a particle was then released from the centre to fly towards the edge of the circle 800 ms later. If the particle landed within the boundaries of the bucket, the bucket would turn green for 500 ms and the participant gained 10 points; else, the bucket turned red for 500 ms and lost 10 points. The number of points accumulated over the task was presented in the top right-hand corner for participants to track their performance. Payment was performance contingent; the more points earned, the higher amount of bonus they received at the end, up to a maximum of 1 USD. Confidence ratings were not incentivized.

On each trial, the particle’s landing location on the circle edge was sampled independently and identically from a Gaussian distribution with SD = 12. As such, the particle landed in the same location with small variations determined by noise. The mean of this distribution did not change until a change-point trial was reached, where it was re-sampled from a uniform distribution U(1,360) (i.e. the number of points on the circle). Participants would therefore have to learn the mean of the new generative distribution after a change-point. The probability of a change-point occurring on each trial was determined by the hazard rate. In the task, there were two hazard rate conditions that varied the number of change-points in a stretch of 150 trials each: stable (hazard rate = 0.025, 4 change-points), and volatile (hazard rate = 0.125, 19 change-points). Hazard rate conditions were not relevant to the analyses of the current paper. The order of hazard rate conditions was randomly shuffled, as were the order of change-points within a condition. Participants completed 300 trials in total, divided into 4 blocks of 75 trials, with no explicit indication when a change in condition block occurred. Breaks were given between blocks which did not fall before the switch of a new hazard rate condition.

Before the start of the task, participants were instructed on the aim of the experiment and shown its layout. Participants then completed 10 practice trials that were excluded from the analysis and did not count for their final score. After the practice, they had to answer 5 questions pertaining to the task. If they answered any of the questions wrong, they would be brought back to the beginning of the instructions and taken through the practice block again. Additionally, in order to reduce the number of participants failing to utilize the confidence scale properly, the task was reset to the beginning if participants left their confidence ratings as the default score for more than 70% of the trials at the 20th and 50th trial mark. They would have to answer the task questions again before proceeding with the task.

### Self-report psychiatric questionnaires & IQ

Participants completed a range of self-report psychiatric assessments after the behavioural task. To enable application of the transdiagnostic analysis with psychiatric dimensions described in previous studies^14,17^, we administered the same nine questionnaires assessing: *Alcohol addiction* using the Alcohol Use Disorder Identification Test (AUDIT)^29^, *Apathy* using the Apathy Evaluation Scale (AES)^30^, *Depression* using the Self-Rating Depression Scale (SDS)^31^, *Eating disorders* using the Eating Attitudes Test (EAT-26)^32^, *Impulsivity* using the Barratt Impulsivity Scale (BIS-10)^33^, *Obsessive-compulsive disorder* (OCD) using the Obsessive-Compulsive Inventory – Revised (OCI-R)^34^, *Trait anxiety* using the trait portion of the State-Trait Anxiety Inventory (STAI)^35^, *Schizotypy* scores using the Short Scales for Measuring Schizotypy^36^, and *Social anxiety* using the Liebowitz Social Anxiety Scale (LSAS)^37^. The order of these self-report assessments administered was fully randomized. Following the questionnaires, participants completed a Computerized Adaptive Task (CAT) based on items similar to that of Raven’s Standard Progressive Matrices (SPM)^38^ to approximate Intelligence Quotient (IQ).

### Medication status

Participants were asked if they were currently taking medication for a mental health issue, and if so, to indicate the name, dosage and duration. 41 (9.38%) participants were currently medicated.

### Transdiagnostic factors

Raw scores on the 209 individual questions that subjects answered from the 9 questionnaires were transformed into factor scores (‘Anxious-Depression’, ‘Compulsive Behaviour and Intrusive Thought’, and ‘Social Withdrawal’), based on weights derived from a larger previous study (N = 1413)^17^ (Figure S7).

### Action-confidence coupling

Regression analyses were conducted using mixed-effects models written in R, version 3.5.1 via RStudio version 1.1.463 (http://cran.us.r-project.org) with the *lme4* package. We examined the coupling between trial-by-trial action updates (*Action*, the absolute difference of bucket position on trial *t* and *t*+1, as the dependent variable) and confidence (*Confidence*, confidence level on trial *t*+1, z-scored within-participant, as the independent variable) with age, gender and IQ as z-scored fixed effects co-variates. Within-subject factors (the intercept and main effect of *Confidence*) were taken as random effects (i.e., allowed to vary across subjects). To test if psychiatric symptom or transdiagnostic dimension severities were associated to changes in action-confidence coupling, the scores were included as z-scored between-subjects predictors in the basic model above.

### Action and confidence

Simple mixed-effects models were used to analyse the basic relationship of psychiatric symptoms/dimensions with *Action* or *Confidence* as dependent variables with the intercept as the random effect, controlled for age, gender and IQ.

### Computation model describing behaviour dynamics

In the task, several evidence sources were available to participants (e.g. new information gained, surprise from unexpected outcomes and uncertainty of their belief) to estimate the mean of the generative distribution in order to position their bucket at where they hope to catch the greatest number of particles. A quasi-optimal Bayesian learning model was used to estimate parameters thought to underlie task dynamics using functions from Vaghi *et al.*^12^. This included *PE^b^* (model prediction error, an index of recent outcomes), *CPP* (change-point probability, a measure representing the belief of a surprising outcome) and *RU* (relative uncertainty, the uncertainty owing to the imprecise estimation of the distribution mean; labelled as *(1−CPP)*(1−MC)* in Vaghi *et al*.^12^).

### Influence of parameters on action and confidence

Regressions were constructed as mixed-effect models with the model parameters (where *PE^b^* is taken as its absolute) and a *Hit* categorical predictor (previous trial was a hit or miss) as within-subject regressors, controlled for age, IQ and gender. For the regression on *Action*, following prior literature^2,11,12,16^, all predictors except *PE^b^* were implemented as interaction terms with *PE^b^*. For *Confidence*, we used a similar regression model but without the interaction term with *PE^b^* and with the regressand and predictors z-scored at participant level. Main effects of the four predictors were correlated with CIT severity, tested with Pearson’s correlation.

For details of the regression equations and computational model, see Supplemental Methods.

## Data Availability

The code and data to reproduce the main analyses are freely available in an Open Science Framework (OSF) repository, at https://osf.io/2z6tw/.

## Acknowledgements

This work is supported by a Postgraduate Ussher fellowship from Trinity College Dublin to T.X.F.S. and a fellowship from MQ: Transforming Mental Health (MQ16IP13) to C.M.G.. The authors thank Dr. Benedetto de Martino for his helpful comments on the manuscript.

## Author Contributions

T.X.F.S. and C.M.G. conceived of and designed the study and analysis plan. T.X.F.S. coded the experiment, collected and analysed the data. T.X.F.S. and C.M.G. wrote the manuscript.

