## Supplementary Information for "Transdiagnostic Phenotyping Reveals a Host of Metacognitive Deficits Implicated in Compulsivity"

for

Supplemental Methods

**Exclusion criteria.** Participants were excluded if they failed any of the following: (i) In the behavioural task, the confidence scale indicator would always start at either 25 or 75 on every trial. Participants who left their confidence rating as the default score for more than 60% of the trials (n > 180 trials) were excluded (N = 42). (ii) The task was also reset from the beginning if confidence ratings were left as the default score for >70% of the first 50 trials (56 participants (9.82%) restarted the task at least once). Those who had their task reset >5 times were excluded (N = 6). (iii) Participants who had more than 50% correlation between the default score and their selected confidence rating were excluded (N = 109). (iv) Participants with a lower mean confidence where the previous trial was correct than incorrect were excluded (N = 66). (v) Participants who incorrectly responded to a “catch” question within the questionnaires: “If you are paying attention to these questions, please select ‘A little’ as your answer” were excluded (N = 16).

**Action-confidence coupling.** First, we measured the coupling between action updates (i.e. the tendency to move the bucket) and confidence. *Action* (the absolute difference of bucket position on trial *t* and *t*+1) was the dependent variable and *Confidence* (confidence level on trial *t*+1) was the independent variable in a trial-by-trial regression analysis with age, gender and IQ as fixed effects co-variates (as with all subsequent analyses). Within-subject factors (the intercept and main effect of *Confidence*) were taken as random effects (i.e., allowed to vary across subjects). *Confidence* was z-scored within-participant, while the fixed effect predictors were z-scored across participant. If action and confidence are appropriately coupled, participants should move the bucket more (larger *Action*) when their confidence levels were low, producing a significant negative main effect of *Confidence* on *Action*. In the syntax of the *lmer* function, the regression was: Action ~ Confidence * (Age + IQ + Gender) + (1 + Confidence | Subject).

We then tested if psychiatric symptom severity was associated to changes in action-confidence coupling by including the total score for each questionnaire (*QuestionnaireScore*, z-scored) as a between-subjects predictor in the model above. Separate regressions were performed for each individual symptom due to high correlations across the different psychiatric questionnaires. The extent to which questionnaire total scores contribute to changes in action-confidence coupling is indicated by the presence of a significant *Confidence***QuestionnaireScore* interaction. A positive interaction effect indicates decreased action-confidence coupling (i.e., decoupling), while a negative interaction effect indicates greater action-confidence coupling. The model was specified as: Action ~ Confidence * (*QuestionnaireScore* + Age + IQ + Gender) + (1 + Confidence | Subject). For the transdiagnostic analysis, we included all three factors in the same model, as correlation across variables was lessened in this formulation and thus more interpretable (only 3 moderately correlated variables r = 0.34 - 0.52, instead of 9 that ranged from *r* = 0.13 - 0.84). We replaced *QuestionnaireScore* in the model formula described previously with three psychiatric factors (*AD, CIT, SW*) entered as z-scored fixed effect predictors. The model was: Action ~ Confidence * (AD + CIT + SW + Age + IQ + Gender) + (1 + Confidence | Subject).

**Action and confidence.** To analyse the basic relationship between task-related variables and psychiatric factors, the analysis approach was the same, but simpler. Dependent variables were: 1) Size of bucket updates (*Action*) and 2) reported confidence (*Confidence*). The models were simply: Task Variable ~ AD + CIT + SW + Age + IQ + Gender + (1 | Subject).

**Computation model describing behaviour dynamics.** In the behavioural task, participants were required to learn the mean of the underlying generative distribution in order to position their bucket at where they hope to catch the greatest number of particles. Their belief on where the particle landing distribution mean could be guided by 1) information gained from the most recent outcome (i.e. moving the bucket with every small shift in particle location), 2) surprising large changes signalling a change in mean distribution (i.e. change-points) and 3) their uncertainty of the distribution mean based on particle landing location experience over trials. To separate these contributions, a quasi-optimal Bayesian computational learning model was used to estimate these parameters thought to underlie task dynamics with MATLAB R2018a (The MathWorks, Natick, MA) using functions from Vaghi *et al.*^1^. (see **Quasi-optimal Bayesian computational model** for further model details). This included *PE^b^* (model prediction error, an index of recent outcomes), *CPP* (probability that a trial was a change-point, a measure representing the belief of a surprising outcome) and *RU* (relative uncertainty, the uncertainty owing to the imprecise estimation of the distribution mean; labelled as *(1-CPP)*(1-MC)* in Vaghi *et al.* (Vaghi *et al.*, 2017)). These parameters (where *PE^b^* is taken as its absolute) together with a *Hit* categorical predictor (previous trial was a hit or miss) were used to regress participant adjustments against the benchmark Bayesian model to investigate participant adjustments in reported confidence (*Confidence*; z-scored confidence level on trial *t*) and bucket movements (*Action*) according to the particle landing locations experienced.

**Influence of parameters on action and confidence.** For the regression on *Action*, following Vaghi *et al.*^1^ and prior literature^2–4^, all predictors except *PE^b^* were implemented as interaction terms with *PE^b^*. For Confidence, we used a similar regression model but without the interaction term with *PE^b^* and with the regressand and predictors z-scored at participant level. Regressions were constructed as mixed-effect models controlled for age, IQ and gender, with the interaction term and main effect of regressors as random effects. The model syntax was written as: Dependent Variable ~ (PE^b^ + CPP + RU + Hit)*(Age + IQ + Gender) + (1 + PE^b^ + CPP + RU + Hit | Subject). We obtained similar regression estimates with Vaghi *et al.*^1^, suggesting that action/confidence is appropriately updated with these parameters describing belief updating (Table S1). The main effects of the four predictors were correlated with CIT severity, where Pearson’s correlation was used to measure the association between symptom dimension severity and the influence of the learning parameters on action update/confidence. To include all three psychiatric factors in the same analysis model (Figure S6), we entered the transdiagnostic factors as additional z-scored fixed effect predictors into the basic model above, where the equation was: Dependent Variable ~ (PE^b^ + CPP + RU + Hit)*(AD + CIT + SW + Age + IQ + Gender) + (1 + PE^b^ + CPP + RU + Hit | Subject). For confidence, a positive interaction between a factor score and *PE^b^, CPP, RU* indicates that higher scores on that factor are associated with a decrease in influence of these parameters on confidence. The converse was applicable for significant *Hit**Factor interactions (as main effect of *Hit* on *Confidence* is opposite signed). For action, as main effect of the parameters on *Action* is inverse from the main effects on *Confidence*, significant parameter*Factor interactions are interpreted in reverse.

**Quasi-optimal Bayesian computational model.** To employ model-based analysis, we calculated task prediction error (PE: distance between the particle landing location and the centre of the bucket) and human learning rate (LR^h^: change in chosen bucket position from t to t+1 divided by PE on trial t) for each trial. Trials where LR^h^ exceeded the 99th percentile (LR^h^ > 7.75) and PE = 0 are thought to be unrelated to error-driven learning^5^, and were thus excluded from analyses with the model parameters (3.05% of total trials).

A learning model was used to determine how participants’ beliefs of the mean of the particle distribution evolved over time. This was estimated by fitting a reduced quasi-optimal Bayesian learner, in accordance with prior literature^1–3,5^, to particle landing location data. Briefly, the model approximates optimal inference by tracking factors that should drive learning: 1) information gained from the most recent outcome (i.e. moving the bucket with every small shift in particle location), 2) surprising large changes signalling a change in mean distribution (i.e. change-points) and 3) the uncertainty of the distribution mean based on particle landing location experience over trials. Using a delta-rule, the model updates its estimate of belief about the particle landing location distribution:

$$B_{t+1}= B_{t}+ \alpha_{t} \times\delta_{t}$$

*B* is the new belief estimate on each trial *t*, which is equal to a point estimation of the mean of the Gaussian distribution where particle locations were sampled (i.e. 1 to 360). Its update is dependent on the learning rate $\alpha$ (LR^b^) and model prediction error $\delta$ (PE^b^). PE^b^ is calculated as the difference between the belief estimate *B_t_* and the new particle landing location *X_t_* and is a measure of information gained from the most recent trial.

$$\delta_{t}= X_{t}- B_{t}$$

As with common reinforcement learning models, LR^b^ determines how much new information (PE^b^) will update the belief estimate. However, LR^b^ is dynamic in the current model i.e. can change on every trial. If LR^b^ = 0, new evidence has had no impact on the update of the belief estimate, while LR^b^ = 1 suggests that the new belief estimate is entirely determined by the most recent outcome. The magnitude of LR^b^ is dependent the statistics of environment with the equation:

$$\alpha_{t}= \Omega_{t}+({1-\Omega}_{t}) ({1-\nu}_{t})$$

The first term, the change-point probability$\Omega$ (CPP), represents an estimate of how likely a change in particle location distribution mean has occurred on a given trial. The second term, model confidence $\nu$, represents the uncertainty due to an inaccurate estimation of the mean. For regression analyses, $(1-\Omega) (1-\nu)$ was labelled as RU (as the additive inverse of MC is relative uncertainty). These two components allow the model to appropriately update belief according to (i) unexpected changes in the environment (change-points) and (ii) the uncertainty about the distribution mean - thus informing when to disregard outliers when the mean is certain. New outcomes are more influential when the model believes that the distribution mean has changed (i.e. CPP is large) or is less sure about the true distribution mean (i.e. MC is small).

The model generates CPP as the relative likelihood that a new particle location is sampled from a new distribution during a change-point (mean determined by a uniform distribution *U* over all 360 possible locations) or drawn from the same Gaussian (*N*) where the current belief estimate *B_t_* is centered upon. These are influenced by the hazard rate *H*, the probability that the mean of the distribution has changed. We set *H* equal to the hazard rates of the task trials (H = 0.025 or 0.125, depending on the trial condition). When the probability of the new particle location coming from a new distribution is high, CPP will be close to 1.

$$\Omega_{t}= \frac{U\left( X_{t} | 1,360 \right)H}{U\left( X_{t} \right|1,360)H+N\left( X_{t} \right|B_{t},\sigma_{t}^{2})(1-H)}$$

$\sigma_{t}^{2}$ is the estimated variance of the predictive distribution, which consists of the variance of the generative Gaussian distribution $\sigma_{N}^{2}$ and the generative variance modulated by MC ($\nu$). As the predictive distribution variance is dependent on MC, it is larger than the generative variance where MC is the smallest (i.e. after change-points, where uncertainty of the new distribution mean is the highest) and will slowly reduce towards the generative variance. Thus, the model describes particle locations occurring after a change-point as less likely sampled from another new distribution.

$$\sigma_{t}^{2}= \sigma_{N}^{2}+ \frac{\left( {1-\nu}_{t} \right)\sigma_{N}^{2}}{\nu_{t}}$$

Lastly, MC is computed for the subsequent trial with a weighted average of the generative variance conditional on a change-point (first term), generative variance conditional on no change-point (second term), and variance due to the model’s uncertainty of whether a change-point occurred (third term) in the numerator. The denominator includes the same terms plus just the generative distribution variance ($\sigma_{N}^{2}$) representing the uncertainty owing to noise. The full equation is as follows:

$$\nu_{t+1}= \frac{\Omega_{t}\sigma_{N}^{2}+\left( {1-\Omega}_{t} \right)\left( {1-\nu}_{t} \right)\sigma_{N}^{2}+\Omega_{t}\left( {1-\Omega}_{t} \right)\left( {\delta_{t}\nu}_{t} \right)^{2}}{\Omega_{t}\sigma_{N}^{2}+\left( {1-\Omega}_{t} \right)\left( {1-\nu}_{t} \right)\sigma_{N}^{2}+\Omega_{t}\left( {1-\Omega}_{t} \right)\left( {\delta_{t}\nu}_{t} \right)^{2}+\sigma_{N}^{2}}$$

Supplemental Figures and Table

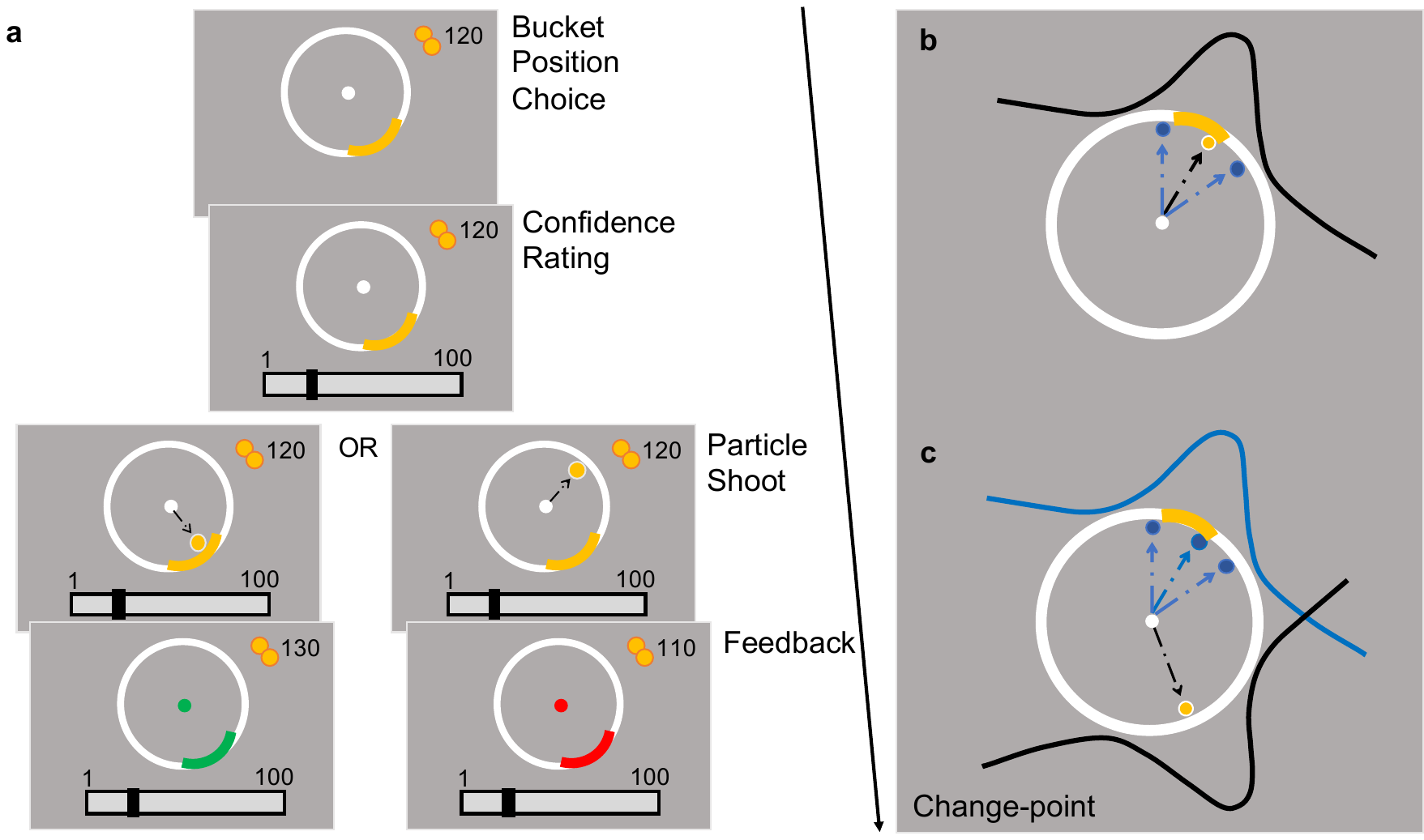

**Figure S1.** Predictive-inference task.

**(a) Trial sequence.** Participants were instructed to position a bucket (yellow arc on the circle edge) to catch a flying particle, and thereafter rated their confidence that they would catch the particle. Particles were fired from the centre of the circle to the edge. Points were gained when the particle was caught, and the bucket turned green; else, points were lost and the bucket turned red.

**(b) Particle trajectories.** For every trial, landing locations were independently sampled from a Gaussian distribution. As such, particles landed around the same area with small variations induced by noise. For illustration purposes, dashed arrow lines represent particle trajectory of current (black) and past (blue) trials, which over trials allow subjects to generate a representation of the Gaussian.

**(c) Change-points.** The mean of the distribution abruptly moves to another point on the circle when a “change-point” occurs. This new mean is then sampled in the same manner as **(b)** until the next change-point.

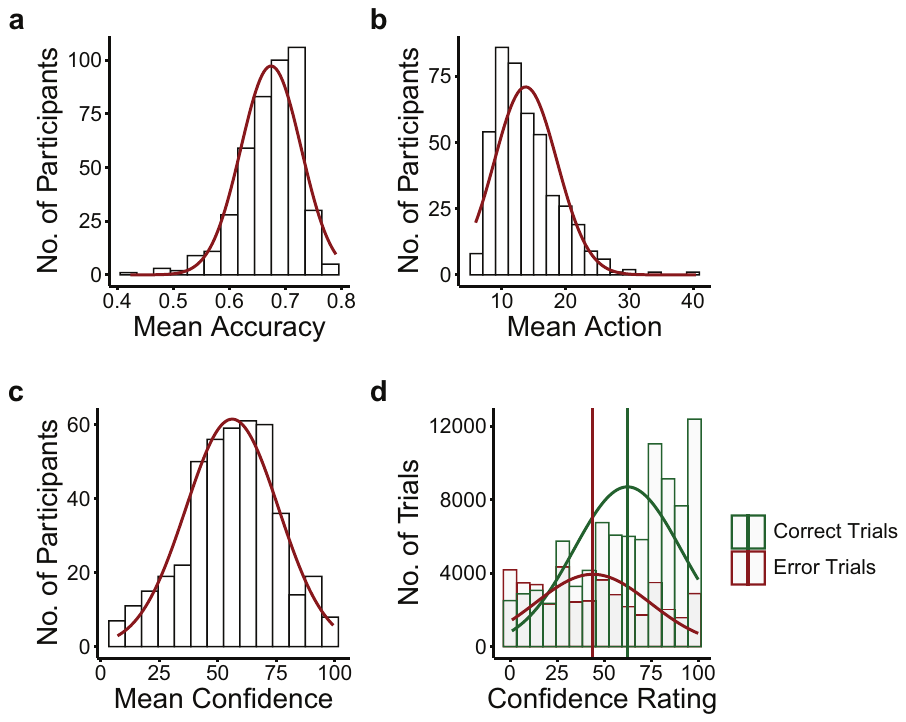

**Figure S2.** Behavioural results. Across participants, the distribution of:

**(a)** Mean accuracy.

**(b)** Mean action (the tendency to move the bucket).

**(c)** Mean confidence level.

**(d)** Confidence ratings for correct (green) and incorrect (red) trials. Vertical lines denote mean confidence level for respective trial type.

Across participants, mean accuracy ranged from 42.33% to 79.00% (mean = 67.42%, SD = 5.38%; Figure S2a), mean action (tendency to move bucket position) ranged from 5.88 to 40.44 (mean = 13.74, SD = 4.91, Figure S2b) and mean confidence level ranged from 7.21 to 99.39 across participants (mean = 56.19, SD = 19.85; Figure S2c). Performance accuracy accounted for only 1.7% of the variance in confidence levels (between-subject correlation: *r* = 0.13, *p* *<* 0.009). Participants were using the confidence scale appropriately, giving higher confidence after correct trials (mean = 62.42, SD = 28.53), and lower confidence after incorrect trials (mean = 43.98, SD = 30.45) (Figure S2d).

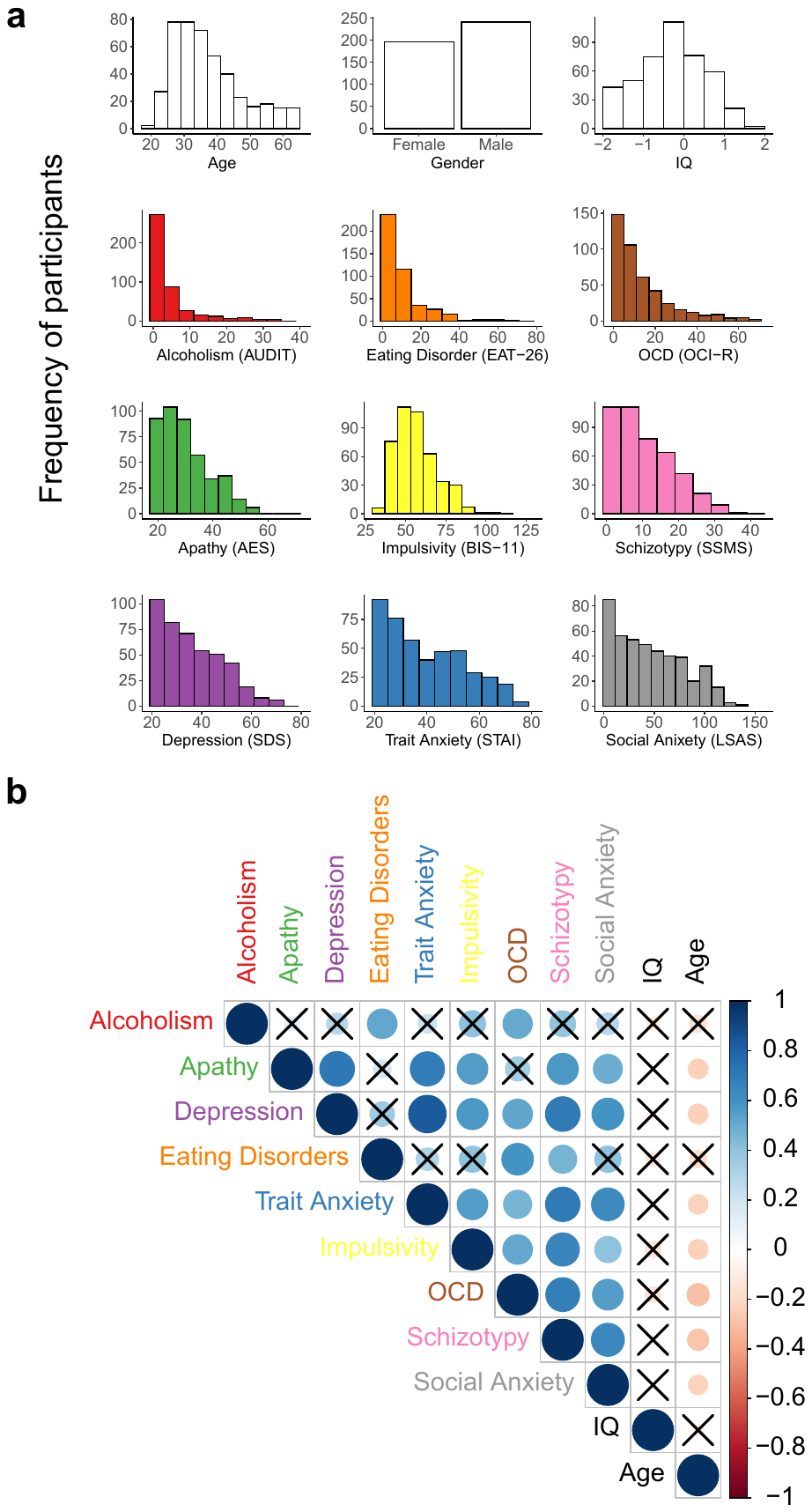

**Figure S3.** Demographics and self-reported psychopathology spread.

**(a)** Age, IQ and psychiatric symptoms score distributions across participants.

**(b)** Correlation matrix of mean scores of the nine questionnaires, age and IQ. Colour scale indicates correlation coefficient, size of colour patch indicates significance. X denotes correlation fails 95% Confidence Interval.

**Table S1.** Effects of Bayesian Model Parameters on Action and Confidence. SE = standard Error, CI = confidence interval.

| **Predictor** | **β (SE)** | **95% CI** | **t-value** | **p-value** |
| --- | --- | --- | --- | --- |
|  | Regression on Action | | | |
| PE^b^ | 0.33 (0.02) | [0.27, 0.38] | 11.61 | < 0.001 *** |
| CPP | 0.46 (0.02) | [0.41, 0.50] | 20.06 | < 0.001 *** |
| RU | 1.37 (0.08) | [1.21, 1.52] | 17.25 | < 0.001 *** |
| Hit | -0.77 (0.02) | [-0.81, -0.73] | -40.54 | < 0.001 *** |
|  | Regression on Confidence | | | |
| PE^b^ | -0.04 (0.01) | [-0.06, -0.02] | -3.57 | < 0.001 *** |
| CPP | -0.20 (0.02) | [-0.24, -0.17] | -12.16 | < 0.001 *** |
| RU | -0.24 (0.01) | [-0.27, -0.21] | -17.15 | < 0.001 *** |
| Hit | 0.26 (0.01) | [0.23, 0.29] | 22.84 | < 0.001 *** |

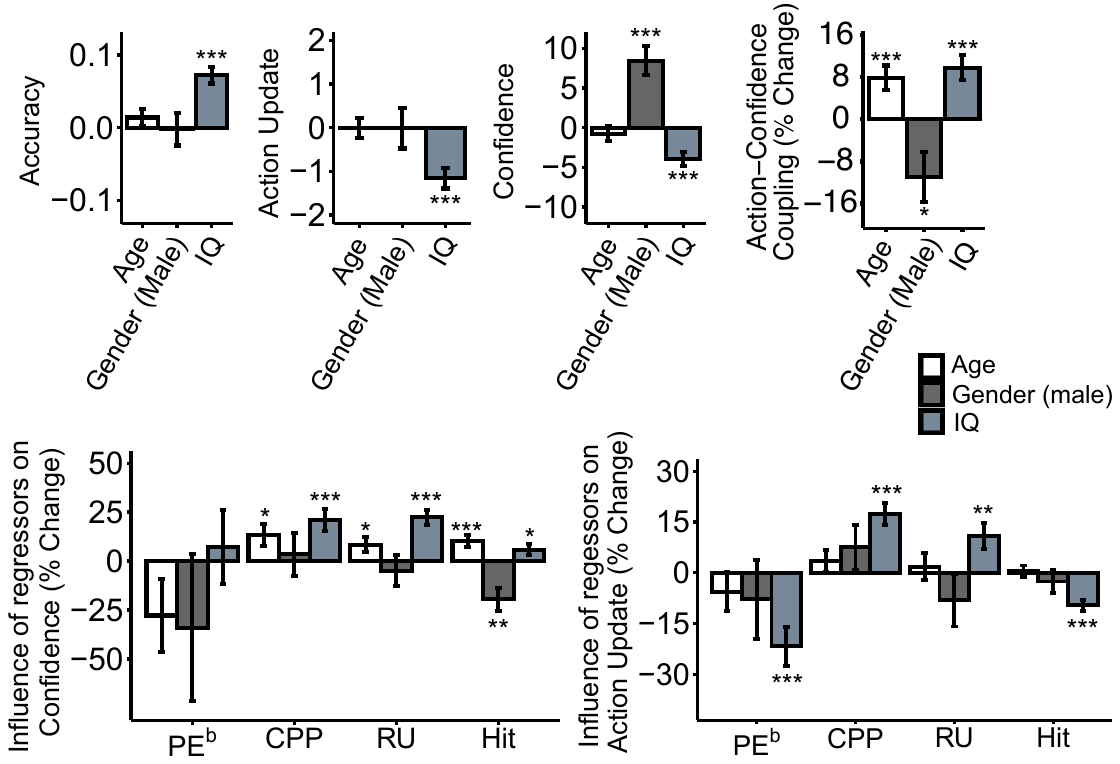

**Figure S4.** Associations between age, gender and IQ with accuracy, action update, reported confidence, action-confidence coupling or the influence of the model predictors (PE^b^, CPP, RU) and Hit on confidence/action update. Error bars denote standard errors. The Y-axes indicates the change/percentage change in each dependent variable as a function of 1 standard deviation increase of demographic scores. *p < 0.05, **p < 0.01, ***p < 0.001.

We tested in an exploratory fashion for relationships of task accuracy, action and confidence with age, IQ and gender (Figure S4). IQ was found to predict better performance (β = 0.07, SE = 0.01, 95% CI [0.05, 0.09], p < 0.001), lower action updating, (β = -1.16, SE = 0.23, 95% CI [-1.62, -0.71], p < 0.001) and lower confidence (β = -3.97, SE = 0.92, 95% CI [-5.77, -2.17], p < 0.001). Additionally, gender (male) was associated with higher confidence (β = 8.43, SE = 1.85, 95% CI [4.81, 12.06], p < 0.001).

**IQ, age and gender were controlled for in all analyses.** Increased action-confidence coupling was associated to age (β = -0.70, SE = 0.21, 95% CI [-1.10, -0.29], p < 0.001), and IQ (β = -0.87, SE = 0.21, 95% CI [-1.27, -0.46], p < 0.001) while decreased in males (β = 0.97, SE = 0.42, 95% CI [0.16, 1.78], p = 0.02). For the model-based trial-wise analyses, age was related to an increased influence of CPP (β = -0.03, SE = 0.01, 95% CI [-0.05, -0.01], p = 0.02), RU (β = -0.02, SE = 0.01, 95% CI [-0.04, -0.002], p = 0.03) and Hit (β = 0.02, SE = 0.01, 95% CI [0.01, 0.04], p = 0.03) on confidence. Males were associated to an increased influence of Hit (β = -0.05, SE = 0.02, 95% CI [-0.08, -0.02], p = 0.001) on confidence, while IQ predicted increased influence of CPP (β = -0.05, SE = 0.01, 95% CI [-0.06, -0.02], p < 0.001), RU (β = -0.05, SE = 0.01, 95% CI [-0.07, -0.03], p < 0.001) and Hit (β = 0.02, SE = 0.01, 95% CI [0.0003, 0.03], p = 0.05) on confidence. For action update, only IQ effects were significant – it was related to an increase in CPP (β = 0.08, SE = 0.02, 95% CI [0.05, 0.11], p < 0.001) and RU (β = 0.15, SE = 0.05, 95% CI [0.04, 0.25], p = 0.006) influence, and decreased PE^b^ (β = -0.07, SE = 0.02, 95% CI [-0.11, -0.03], p < 0.001) and Hit (β = 0.07, SE = 0.01, 95% CI [0.05, 0.10], p < 0.001) influence on action update.

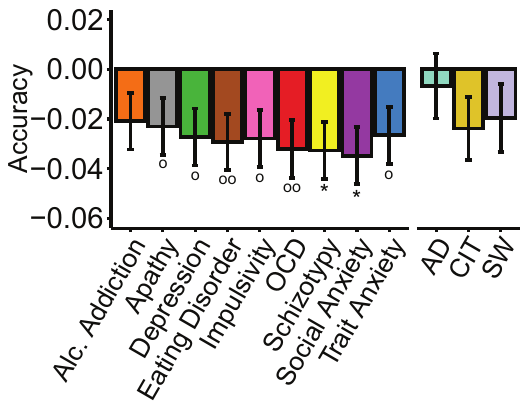

**Figure S5**. Associations between accuracy (hit (1) or miss (0)) with psychiatric symptoms or transdiagnostic factors, controlled for age, IQ and gender. Error bars denote standard errors. The Y-axis indicates the change in accuracy as a function of 1 standard deviation of symptom/factor scores. ^o^p < 0.05, ^oo^p < 0.01 uncorrected, *p < 0.05. Results are Bonferroni corrected for multiple comparisons over number of symptoms/factors.

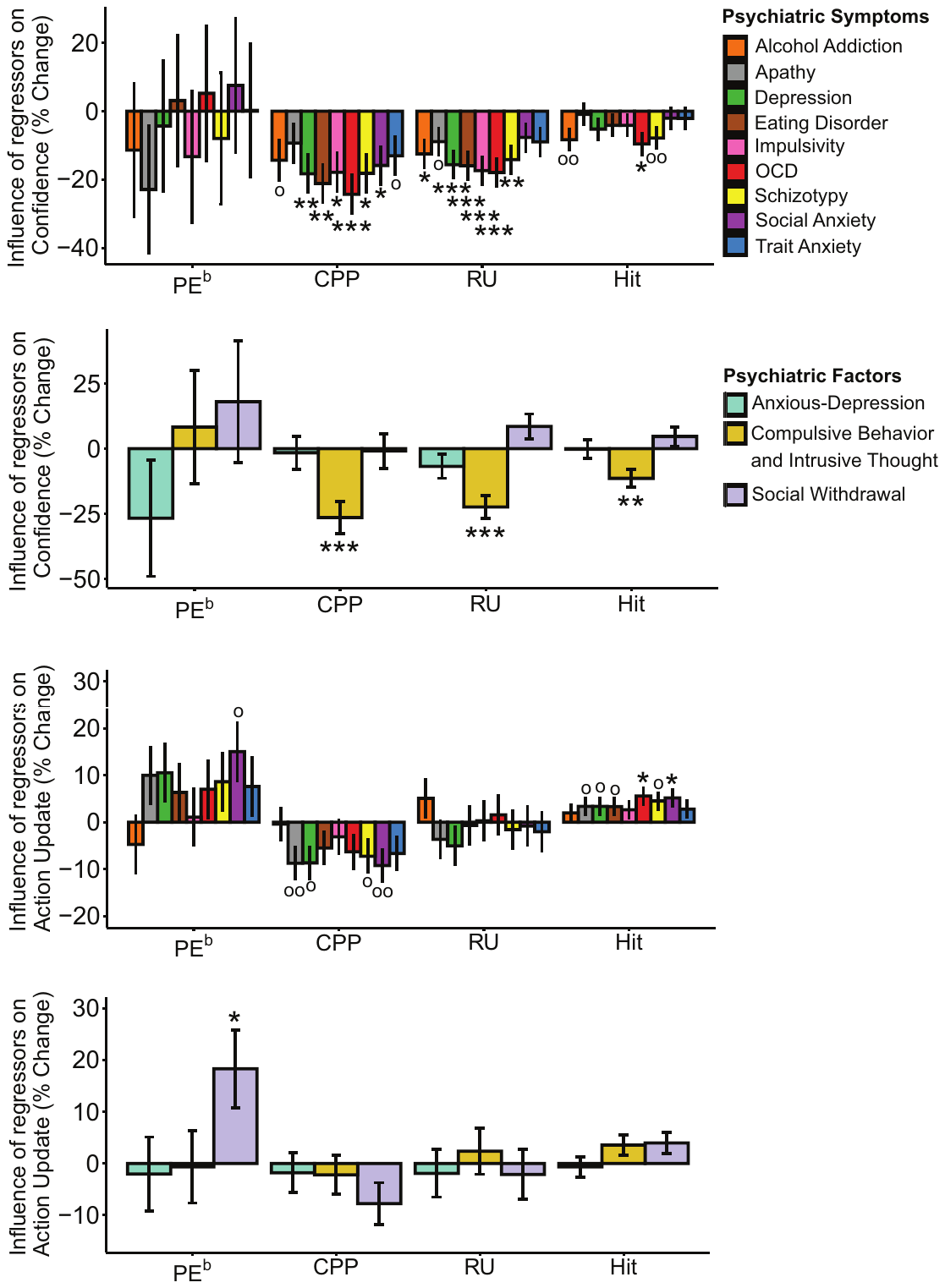

**Figure S6**. Regression analyses of trial-wise confidence and action adjustments with psychiatric symptoms/factors, controlled for age, IQ and gender. Each psychiatric symptom was examined in a separate regression, whereas factors were included in the same model. Error bars denote standard errors. The Y-axes indicate the percentage change in each dependent variable as a function of 1 standard deviation of symptom/factor scores. ^o^p < 0.05, ^oo^p < 0.01 uncorrected, *p < 0.05, ***p < 0.001. Results are Bonferroni corrected for multiple comparisons over number of symptoms/factors.

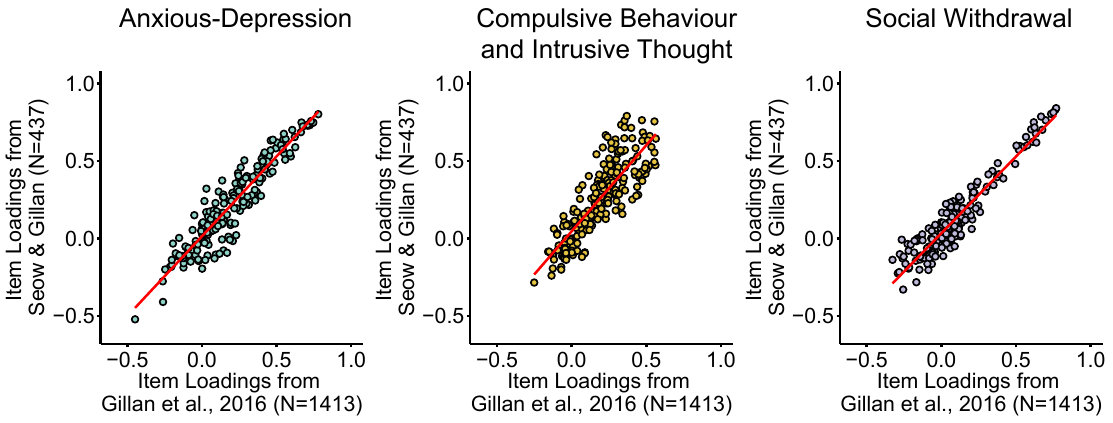

**Figure S7**. Correlations between item loadings obtained from the factor analysis in Gillan et al. (2016) and the present study for each psychiatric symptom dimension. Questionnaire item loadings were highly correlated for all three factors (Anxious-depression: r = 0.94; Compulsive behaviour and intrusive thought: r = 0.85, Social withdrawal: r = 0.95), supporting the reproducibility of the psychiatric symptom dimensions.

**Transdiagnostic symptom dimensions are reproducible.** Transdiagnostic factors scores (‘Anxious-depression’, ‘Compulsive behaviour and intrusive thoughts’, ‘Social withdrawal’) in the present study were derived from weights obtained from a prior larger study (N = 1413)^6^. This 3-factor structure was previously reproduced in a smaller independent sample (N = 497)^7^, and here we again replicated similar psychiatric dimensions with our current data (N = 437) with the factor analysis (Figure S7). For further details of the factor analysis methodology, see Gillan et al.^6^.

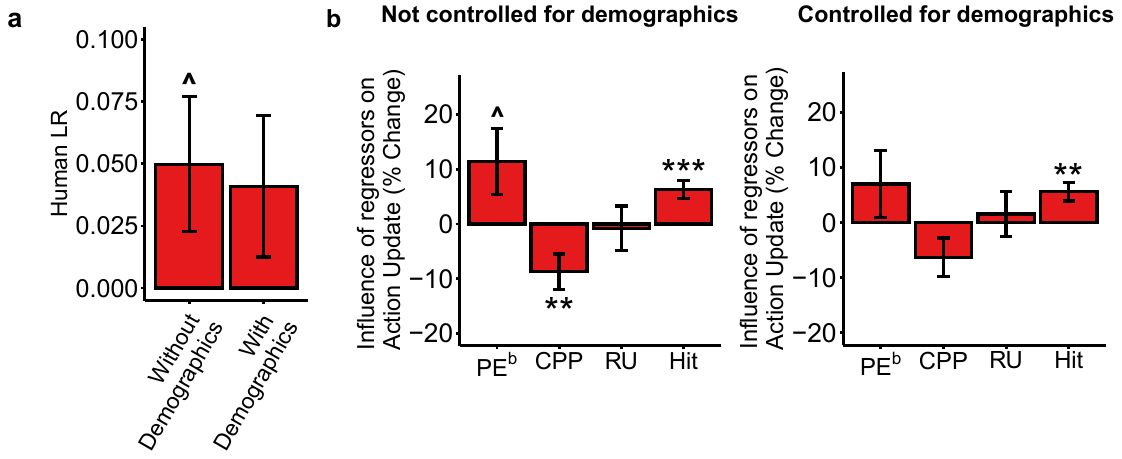

**Figure S8**. Regression analyses of **(a)** human learning rate (ratio of bucket movement and task prediction error) and **(b)** action adjustments in OCD, in a model that controlled for age, IQ and gender and in a model that did not. Error bars denote standard errors. The Y-axes indicate the change/percentage change in dependent variable as a function of 1 standard deviation of OCD symptom scores. ^^^p < 0.07, **p < 0.01, ***p < 0.001. Results are not Bonferroni corrected for multiple comparisons.

**Action updating effects in OCD with/without controlling for demographics.** Vaghi *et al.*^1^ reported that OCD patients exhibited a higher mean learning rate and that their action updates were more strongly influenced by recent information (PE^b^) and less to large unexpected environmental changes (CPP). In the course of exploring the source of this discrepancy with our data, we found that when we repeated our analysis without controlling for age, gender and IQ, some of their effects were recovered here. OCD symptoms were associated with changes in learning and sensitivity to both PE^b^ and CPP in action updating. Specifically, LR^h^ (human learning rate) (*β* = 0.05, *SE* = 0.03, 95% CI [-0.003, 0.10], *p* = 0.07, uncorr.) and the influence of PE^b^ on action showed a trend towards a positive association with OCD symptoms (*β* = 0.04, *SE* = 0.02, 95% CI [-0.001, 0.07], *p* = 0.06, uncorr.) and the influence of CPP on action showed a negative association with OCD symptoms (*β* = -0.04, *SE* = 0.02, 95% CI [-0.07, -0.01], *p* = 0.007, uncorr.). These discrepancies suggest that demographic characteristics perhaps partially explain the pattern of action updating effects observed in the prior patient study (Figure S8).

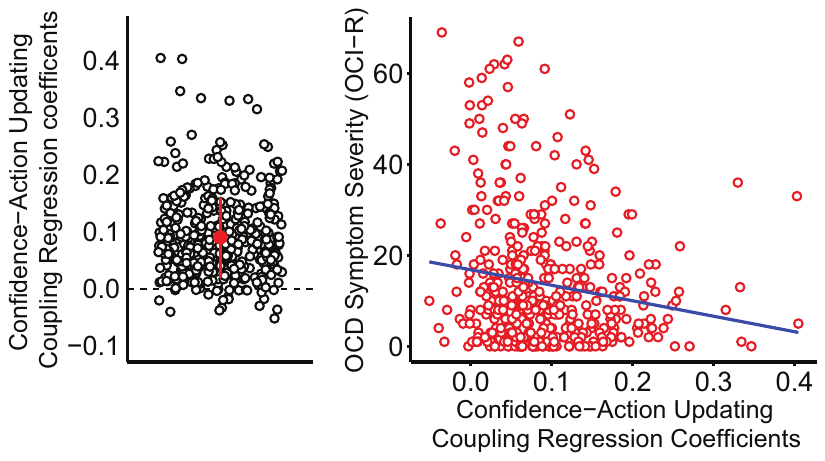
 **Figure S9.** Regression model where confidence updating was predicted by action updating. Dots represent coefficient estimates for individual participants. Red marker indicates mean and SD. These coefficients were correlated with OCD symptom severity, where confidence-action updating coupling was observed to decrease with increasing OCD symptom severity (r = -0.18, p < 0.001).

**Action-confidence decoupling analysis.** Although this has no bearing on our results (or theirs), we note that Vaghi *et al.*^1^ defined action-confidence coupling slightly differently to how we chose to define it in the present paper – they used confidence *updating* (i.e. absolute difference between z-scored confidence from trial *t* and *t*-1), instead of the reported confidence level on trial *t*. We suggest that z-scored confidence ratings (rather than their change from trial to trial) are more appropriate because this accounts better for instances where a person has several relatively large PEs in a row (as they figure out where to place the bucket), and should thus not rationally ‘change’ their confidence rating in response to these PEs, but maintain it at a low level. Although we flag this for the interested reader, we underscore that the two measures are correlated and indeed when we use their definition, we similarly show that self-reported OCD symptom severity predicts confidence-action updating decoupling (*r* = -0.17, *t* = -3.58, 95% CI [-0.26, -0.07], *p* < 0.001, Figure S9).
